# Mosaic Mutations in Blood DNA Sequence Are Associated with Solid Tumor Cancers

**DOI:** 10.1101/065821

**Authors:** Mykyta Artomov, Manuel A. Rivas, Giulio Genovese, Mark J. Daly

**Affiliations:** Broad Institute, Cambridge, MA, USA, 02139; Analytic and Translational Genetics Unit, MGH, Boston, MA, USA, 02114; Department of Chemistry and Chemical Biology, Harvard University, Cambridge, MA, USA, 02138

## Abstract

Recent findings in understanding the causal role of blood-detectable somatic protein-truncating DNA variants in leukemia prompt questions about generalizability of such observations for other cancer types. We used exome sequencing to compare 22 different cancer phenotypes from TCGA data (~8,000 samples) with more than 6,000 controls using a case-control study design and demonstrate that mosaic protein truncating variants in these genes are also associated with solid-tumor cancers. We analyzed tumor DNA samples from TCGA and observed that the cancer-associated mosaic variants are absent from the tumors.

Through analysis of different cancer phenotypes we observe gene-specificity for mosaic mutations. PPM1D in previous reports has been linked to breast and ovarian cancer, which our analysis confirms as a specifically associated to ovarian cancer. Additionally, glioblastoma, melanoma and lung cancers show gene specific burden of the mosaic protein truncating mutations. Taken together, these results extend existing observations broadly and link solid-tumor cancers to somatic blood DNA changes.

## Introduction

Several recent studies^1, 2, 3^ have reported associations of mosaic protein truncating variants (PTV) in *PPM1D, TET2, ASXL1* and *DNMT3A* with blood cancers. Intriguingly, such mosaic mutations in *PPM1D* have also been convincingly associated with breast and ovarian cancer - however, since these mutations are somatic, rather than germline, a role in causation has not been clear. We sought to more fully explore the relationship of these somatic mutations, clearly causally linked to blood cancers, in solid tumor cancer using a large assembly of germline and somatic exome DNA sequences of 7,979 cancer cases from TCGA^4^ and performed a large-scale case-control study with 6,177 population controls with no cancer phenotype reported.

## Results

Using data available from dbGAP, we performed a large-scale joint variant calling of germline DNA samples from the blood of cancer cases and controls - primarily from an assembly of TCGA samples (cases) compared with unselected population controls (with no known cancer status) from several studies (NHLBI-ESP, 1000 Genomes, ATVB, T2D, Ottawa Heart) appropriately consented for broad use as controls. Importantly, all cases and controls in this analysis have age at DNA sampling available (Supp. Table 1).

Observations of the mosaic mutations are very likely affected by several parameters - both biological (age, smoking) and technical (depth of coverage, variant calling accuracy). To make the case-control comparison robust we first identified what adjustments to the model of association are needed. We observed 348 PTVs (stop gain, essential splice site, frameshift mutations) in the four established somatic leukemia genes. Detection of somatic mutations with low non-reference allele balance depends importantly on depth. In order to insure no different sensitivity in our cases and controls we first compared depth of coverage in these genes in our cancer germline (average 33X coverage) and control (average 29X coverage) data. We further looked specifically at cases and controls with called PTVs. For germline heterozygous sites, the expected allele balance is 0.5 so we applied a binomial test to determine low allele balance genotypes based on the depth of coverage and number of alternative reads. Those with p<0.001 (i.e., heterozygotes with significantly less than 50% non-reference allele) and more than 20x coverage were determined to be mosaic and kept for further analysis (Supp. Fig 1,2). To further investigate any statistical bias due to a coverage of cases and controls - we tested whether there is a statistical difference in coverage and ref/alt reads counts between cancer cases and controls that carry at least one PTV in the 4 candidate genes with generalized linear model testing. The cancer status of the sample appears to be a non-significant (p=0.279, p=0.898 if adjusted for age) parameter, confirming that called PTVs are adequately covered in both cases and controls and protein-truncating mosaic events have equal chances to be detected in both cohorts. We finally evaluated the probability of calling a protein truncating DNA variant in cases and controls with respect to coverage (Sup. Fig. 3), since there is slightly higher sensitivity for the detection of DNA variants in cases we adjusted further analysis for the coverage differences. From these analyses, we conclude that all minor technical differences in sensitivity to find mosaic variants in cases and controls were accounted for - a pre-requisite for subsequent analyses.

We then assessed association between mosaic PTV and cancer status by generating a data set consisting of 7,979 cancer cases and 6,177 controls (See Methods). We applied a binomial generalized linear model considering age, coverage depth and mosaic PTV carrier status and found significant evidence of association with cancer status (P=0.00108, OR=1.26; OR CI=1.1-1.47). Since it was previously shown that *PPM1D* PTVs are associated with breast and ovarian cancers, we removed breast and ovarian cancer samples and repeated analysis, which confirmed the association (P=5.67×10^−4^, OR=1.3; OR CI=1.12-1.52) - suggesting the reported observations regarding *PPM1D* and breast and ovarian cancers are more general. As our set of controls was on average roughly 10 years younger than our cancer cohort and age has been shown to be a strong predictor of the existence of these somatic mosaic events, the inclusion of age in the above model is critical. We further evaluated the effect of the age on the probability of finding a mosaic event (Sup. Fig. 4)^5^ and the generalized linear modeling above was adjusted for age differences between cases and controls. We also adjusted our model for minor coverage differences between cases and controls.

It is known that specifically PTVs in the last exon of *PPM1D* are enriched in cases of breast and ovarian cancer^1^. We observed the same enrichment in our dataset - 18 mosaic PTVs in *PPM1D*, 17 of which appeared in the last exon of the gene. Thus we tested other candidate genes for distribution of the PTVs. Variants in *TET2* also show strong exon specificity - 44 out of 50 total PTVs are found in the 3^rd^ exon of the gene. Out of 40 ASXL1 PTVs 35 appear in the last exon while *DNMT3A* PTVs do not show any exon specificity (Supp Fig. 5). Previous reports of leukemia studies observe accumulation of the mosaic missense mutations in the last exons of *DNMT3A.* We found the same to be true in blood for solid tumor cancer cases as well (p=4.78×10^−4^, Supp Fig. 6).

As it was previously demonstrated, mosaic PTVs in the list of candidate genes have a high association with the development of leukemia as mutations in these genes were demonstrated to precede and predict the development of leukemia, indicating a causative role in leukemias^2, 3, 6, 7, 8^. We next sought a key piece of evidence for evaluating the role of these mutations in solid tumors. Specifically we evaluated the quantity of these mosaic PTVs between tumor and germline DNA across all cancer samples included in this study with a detectable event in blood DNA and observed that mosaic PTVs in the candidate genes present in blood DNA were largely absent in the tumor DNA samples from the same individual (Fig. 1). This strongly indicates that these events in the blood did not represent residual evidence from driver mutations involved in tumor development (in which case we would have expected higher, or perhaps 100% of the mutated allele to be found). As before, we compared coverage in tumor and germline DNA samples, which shows, consistent with the design of TCGA, that tumors have similar or better coverage indicating that the deficit of these mosaic events in tumors is not sensitivity based (Supp. Fig 7). This observation is consistent with the findings of mosaic *PPM1D* variants in breast/ovarian cancers^1^.

**Fig. 1.**
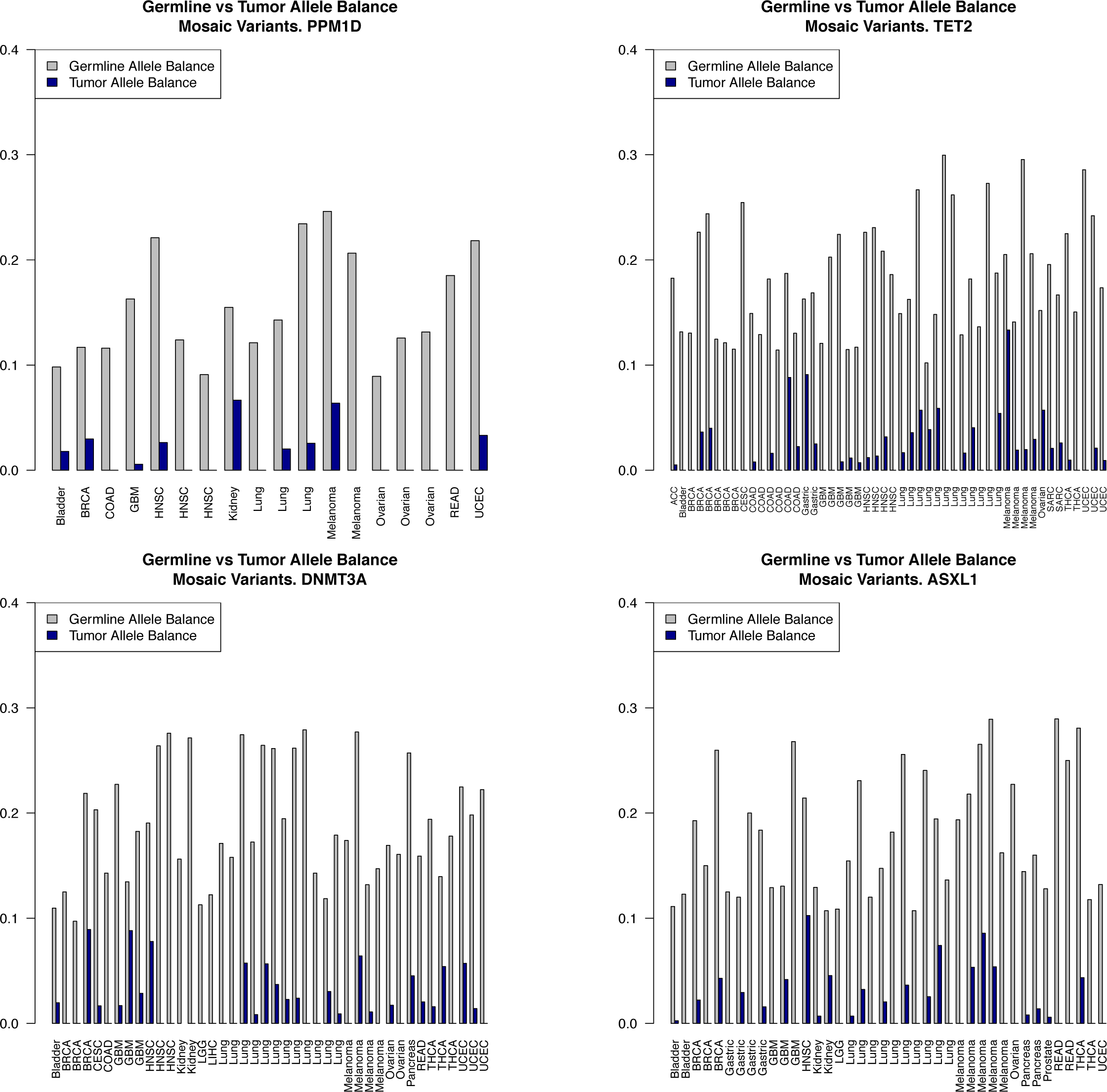
Blood vs Tumor allele balance for each sample with mosaic PTV in (A) PPM1D, (B) TET2, (C) DNMT3A, (D) ASXL1. Observed in blood mosaic mutations are strongly depleted from the tumor somatic DNA (Wilcoxon test P<10^−16^).

We considered whether presence of mosaic PTVs showed any evidence of cancer specificity. Under the null model, we would expect mosaic events to be found in all candidate genes at the same rate in each of the 20 cancer phenotypes. We tested for deviation from this null model and applied a multiple hypothesis testing correction procedure (p=.05/20) to reject the null that the rate is the same across all the phenotypes in all the genes. We first tested if any of the cancer phenotypes shows unusual burden of the mosaic PTVs by randomly drawing sample sets (controlling for similarity of age distributions between the random and target sets) from the total cancer samples cohort and estimating how likely is observed amount of mosaic PTVs in each cancer phenotype. (Fig. 2a). Glioblastoma, melanoma and lung cancers demonstrate significantly increased burden of mosaic PTVs compared to other cancers. We then examined the distribution of mosaic PTVs across the candidate genes in each cancer phenotype (Fig. 2b, Sup. Table 2). Using the same permutation analysis, it appears that several cancer types show a trend for accumulation of the mosaic mutations in specific genes. Intriguingly, ovarian cancer is specifically associated with *PPM1D* mutations, which is supported by the previous report^1^. We also observe associations of head and neck squamous cell carcinoma with *PPM1D*, colorectal adenocarcinoma and glioblastoma with *TET2*. Interestingly, cutaneous melanoma is associated with *ASXL1* mosaic mutations as *ASXL1* has protein-interaction with *BAP1,* a well-established risk factor for melanoma^9^. Lung cancer shows burden of the mosaic mutations that is distributed across several genes, suggesting no specificity in accumulation of the mosaic mutations. This perhaps could be related to the observation that smokers have higher rate of mosaic PTVs.

**Fig. 2.**
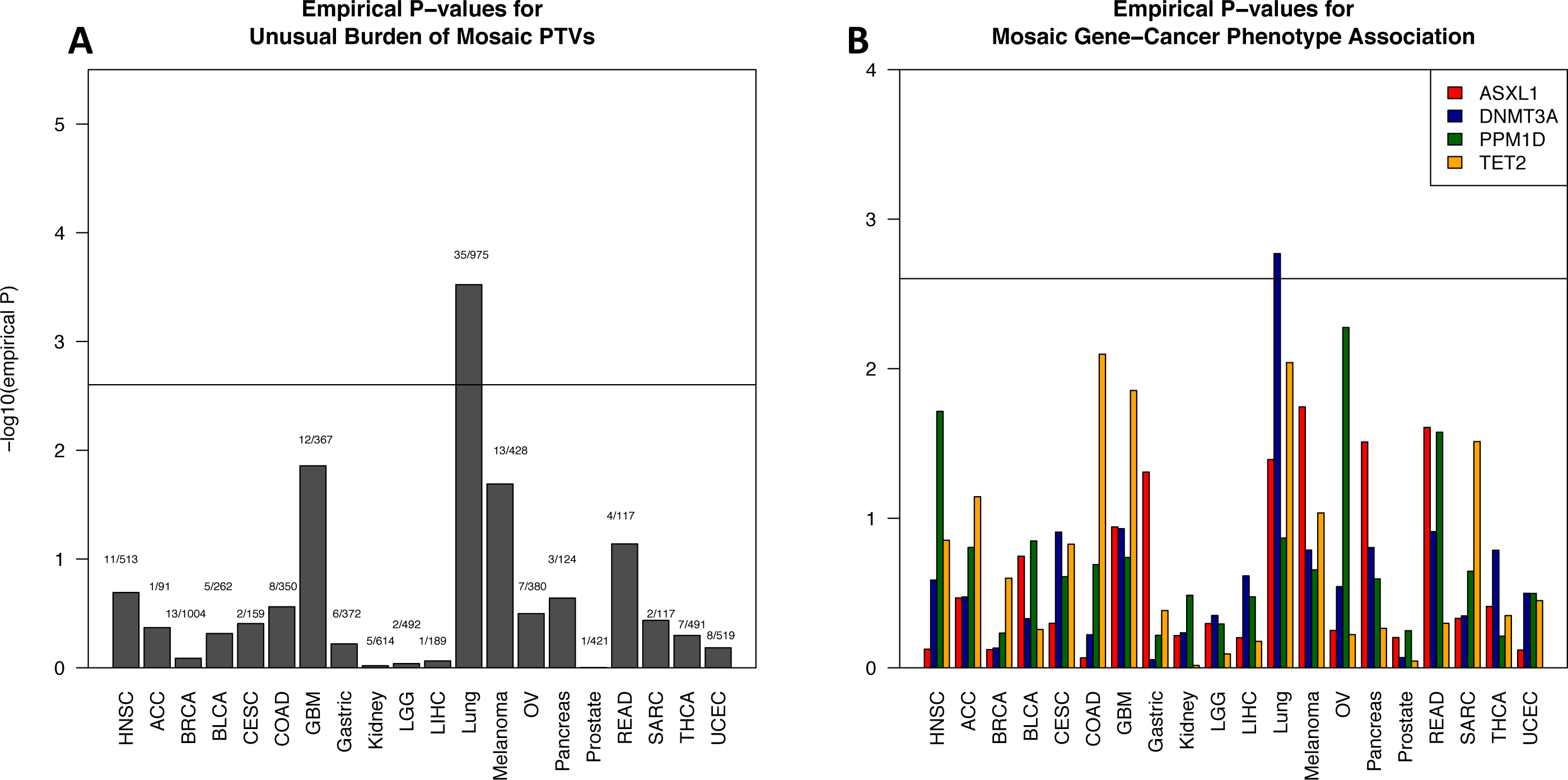
Solid-tumor cancer phenotypes show gene specificity with respect to mosaic PTVs. (A) Empirical enrichment of the different cancer phenotypes with mosaic PTVs (B) Per gene significance of mosaic PTV burden in each cancer phenotype. Experiment-wise significance level is set with Bonferroni correction for multiple phenotypes tested. Ovarian cancer shows previously reported specific association to PPM1D mosaic PTVs.

We used the previously reported set of samples from Swedish national patient registers^2^ to estimate the frequency of mosaic PTVs and associated solid-tumor cancer development in a population unselected for cancer.

We removed from analysis all samples that had an evidence of leukemia or lymphoma developed before the DNA collection as well as those samples that have mosaic missense mutations in *DNMT3A* to estimate the contribution of the PTVs only. The final dataset for this analysis consisted of (92 mosaic PTV carriers and 11,000 non-carriers) samples. There were 15 individuals with pre-DNA collection record of the solid-tumor cancer in the cohort of mosaic PTV carriers and 1,149 samples with record of solid-tumor cancer among non-carriers. We tried using different thresholds for age of the samples to estimate significance of enrichment. However, due to a small incidence of the mosaic mutations in the population unselected for cancer, this test was inconclusive (Supp. Table 3a).

We added mosaic missense *DNMT3A* mutations carriers to the mosaic samples cohort and repeated population analysis (Supp. Table 3b). This resulted in a total of 180 mosaic samples. There were 31 individuals (~17%) with pre-DNA collection record of the solid-tumor cancer in the cohort of mosaic PTV carriers (1,118 (~10%)cancer records in 10,912 non-mosaic samples). Once corrected for age this enrichment appears to be insignificant, thus for samples unselected for cancer a much larger cohort is needed to reach a significant conclusion.

Alternative possible driver of previously reported leukemia association with mosaic PTVs is a clinical intervention, specifically - radiation treatment, well known to be leading to increased risk of leukemia. Within the limited available clinical data in TCGA we saw no clear associations to treatment history (neoadjuvant treatment history (p=0.116), radiation therapy (p=0.348), pathologic tumor stage (p=0.354) or other outcome variables when adjusted for age and cancer subtype with mosaic PTV carrier status (Supp. Tables 4,5,6). However, these analyses are extremely power-limited at this point and, as discussed below, recent evidence supports the idea that, instead of a causal role in promoting solid tumors, these variants are most likely enriched in incidence or survival by chemotherapy or radiation treatment.

## Discussion

Our study investigates association of the mosaic protein-truncating variants in 4 previously associated with blood cancer risk genes with solid-tumor cancer phenotypes.

Previously observed strong association of mosaic PTVs with increased risk of leukemia in our observations is extended to the solid-tumor cancers. There are several possible explanations for such an observation. Recent findings in ovarian and breast cancer suggest a significant role of chemotherapy exposure in observed burden of mosaic PTVs in *PPM1D*^10, 11^. Though our study lacks sufficiently detailed records of chemotherapy treatment to extend those observations, the breadth and robustness of the results here suggest that such effect of treatment exposure may more generally apply to other candidate genes and other cancer phenotype. At the same time observed differences in PTV burden gene specificity according to cancer phenotype suggests that there could be some level of specificity of chemotherapy drugs to cause expansion/survival of certain mutated peripheral blood mononuclear cells clones. Importantly, however, such a link may provide a more general and detectable connection between early solid tumor diagnoses and enriched later incidence of leukemia. Other possible explanations for the observed association could be first, immune system changes in response to early pre-clinical stage of cancer. Our additional screening of early onset cancer cases (breast and ovarian cohort with cancer onset before 35, N=374) shows no enrichment in mosaic PTVs suggesting that this hypothesis is likely irrelevant and age of the samples plays important role (or serving as a trigger) for emergence of clonal expansion. Second, is the causal relationship. While a direct role as tumor drivers is ruled out by the absence of PTVs in tumors, we cannot completely eliminate the possibility that these represent a background cancer risk state but find no strong support for this hypothesis. Given fewer than 1% of the population carries a PTV in one of these candidate genes, a large-scale population study with a long-term pre- and post-cancer DNA collection and detailed treatment details will be needed to confidently answer the question whether blood mosaic PTVs are precursors or result of treatment for solid-tumor cancers.

## Methods

### Dataset

Genotypes dataset was created by joint variant calling of cancer cases and non-cancer controls using HaplotypeCaller (GATK-3.0)^12, 13, 14^ with Broad Institute calling pipeline. For functional annotation of variants we used Variant Effect Predictor by Ensembl^15^

PCA was performed to keep for analysis only samples of European ancestry to eliminate possible population effects. PCA was performed with EIGENSTRAT^16, 17^

Resulting genotype file was used to create a PLINK/SEQ^18^ project for further manipulations.

### Clinical data

For testing relevance of the mosaic PTVs to medical treatment/outcome clinical data was downloaded from TCGA web-site https://tcga-data.nci.nih.gov/tcga/dataAccessMatrix.htm.

### Generalized linear model and statistical tests

For further statistical tests we used R-3.0^19^

---

**Supplementary Fig. 1.** Allele balance for all PTVs with >20X coverage in 4 candidate genes in TCGA cancer samples.

**Supplementary Fig. 2.** Allele balance for all PTVs with >20X coverage in 4 candidate genes in control samples.

**Supplementary Fig. 3.** Probability of observation a protein-truncating variant in cancer and control samples with respect to the coverage. Controls show better or equal chances of PTV detection.

**Supplementary Fig. 4.** Mosaic PTV emergence in blood is strongly correlated with age, however probability of finding such mutations in cancer cases is much greater than in samples with no known cancer history.

**Supplementary Fig. 5.** PTVs in *ASXL1, TET2, PPM1D* show exon specificity.

**Supplementary Fig. 6.** Comparison of coverage between the TCGA blood and TCGA somatic DNA samples. On average tumor DNA has equal or better coverage, than blood DNA.

